# Niche construction in evolutionary theory: the construction of an academic niche?

**DOI:** 10.1101/109793

**Authors:** Manan Gupta, N. G. Prasad, Sutirth Dey, Amitabh Joshi, T. N. C. Vidya

## Abstract

In recent years, fairly far-reaching claims have been repeatedly made about how niche construction, the modification by organisms of their environment, and that of other organisms, represents a vastly neglected phenomenon in ecological and evolutionary thought. The proponents of this view claim that the niche construction perspective greatly expands the scope of standard evolutionary theory and that niche construction deserves to be treated as a significant evolutionary process in its own right, almost at par with natural selection. Claims have also been advanced about how niche construction theory represents a substantial extension to, and re-orientation of, standard evolutionary theory, which is criticized as being narrowly gene-centric and ignoring the rich complexity and reciprocity of organism-environment interactions. We examine these claims in some detail and show that they do not stand up to scrutiny. We suggest that the manner in which niche construction theory is sought to be pushed in the literature is better viewed as an exercise in academic niche construction whereby, through incessant repetition of largely untenable claims, and the deployment of rhetorically appealing but logically dubious analogies, a receptive climate for a certain sub-discipline is sought to be manufactured within the scientific community. We see this as an unfortunate, but perhaps inevitable, nascent post-truth tendency within science.

## Introduction

In recent decades, the phenomenon of niche construction (henceforth, NC) (Odling-Smee 1988) has been receiving a lot of attention in the evolutionary biology literature, with many arguments that a full consideration of this phenomenon as an evolutionary mechanism is a critical and consequential aspect of a new, extended, evolutionary synthesis (Laland 2015; Laland et al. 2015). It has been suggested that an NC perspective goes beyond the standard thinking in evolutionary biology in significant ways, and that an incorporation of an NC perspective necessitates a major overhaul of how we think about the evolutionary process, especially the role of natural selection in promoting the evolution of adaptations (Laland 2015; Laland et al. 2014a, Laland et al. 2014b, Laland et al. 2015). Just to cite a few representative examples from the last four years alone, it has been claimed that:

- “Niche construction theory is more than just an alternative perspective; it is a serious body of formal evolutionary theory” (Laland et al. 2014a);
- “…developmental bias and niche construction may be viewed as essentially the same phenomena expressed inside and outside the organism” (Laland et al. 2014b);
- “Niche construction theory explicitly recognizes environmental modification by organisms (“niche construction”) and their legacy over time (“ecological inheritance”) to be evolutionary processes in their own right” (Odling-Smee et al. 2013); and
- “An extensive body of formal theory explores the evolutionary consequences of niche construction and its ramifications for evolutionary biology and ecology” (Laland et al. 2016).

These are non-trivial claims and, if justified, would certainly indicate that a major rethink of basic concepts in evolutionary biology is in order. In this article, we examine these claims in some detail in order to assess whether these claims are, indeed, justified. As we will argue, a careful examination of these claims suggests that NCT is not quite as consequential as its proponents make it out to be and that, in particular, it does not necessitate any major rethinking of the conceptual structure of evolutionary theory. Several of the points we raise in our critical appraisal of the claims for NCT have been highlighted before (e.g. Dawkins 2004; Brodie III 2005; Wallach 2016), while others are, to our knowledge, novel. However, given how frequently the claims for NCT are repeated, it is perhaps worth repeating the critiques of NCT, too. Otherwise, we risk the possibility of NCT becoming more important in the perception of evolutionary biologists than its reality would justify, in an ironic example of reproductive fitness of an over-blown idea driving an increase in its frequency to the detriment of other, more logically balanced, concepts. We suggest that standard (neo-Darwinian) evolutionary theory (henceforth, SET), properly understood and applied, provides an excellent and parsimonious framework for comprehending the evolutionary process, and that the proponents of NCT are unnecessarily pushing for a far more cumbersome and muddled conceptual construct.

NC is defined as any activity of an organism which modifies its own selective environment, or the selective environments of other, con-or hetero-specific, organisms (Odling-Smee et al. 1996, 2003, 2013). An oft-cited example of NC is the burrowing activity of earthworms in soil that, in turn, alters the morphology and chemistry of the soil by facilitating microbial activity (Hayes 1983; Lee 1985). This process feeds back to future generations of earthworms and places novel selection pressures on their morphological and physiological phenotypes. As pointed out by Wray *et al*. (in Laland et al. 2014c), Darwin’s (1881) last book was on earthworms and dealt, in part, with how “earthworms are adapted to thrive in an environment that they modify through their own activities”. Surprisingly, although ideas about organisms changing their own selective environment have repeatedly been put forward in the past (Darwin 1859, 1881; Fisher 1930; Van Valen 1973, to cite just a few references), modern NCT traces its genesis to a formulation by Lewontin (Odling-Smee et al. 2003), who – in our opinion, rather unfairly and overly simplistically – characterised the SET view of the relationship between organism and environment by using two asymmetrically coupled differential equations (Lewontin 1983, 2000):

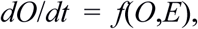

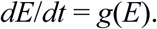

These equations suggest that change in the organismal domain is a function of both organism (*O*) and environment (*E*), whereas change in the environmental domain is solely as a function of the environment itself. Lewontin (1983, 2000) proposed that the implicit assumption that organisms do not change their environments was unjustified and that, therefore, the second equation should be replaced by the following:

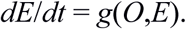

This new system of equations, thus, becomes symmetrically coupled by acknowledging the contribution of organisms to environmental change. This idea was taken further and explored in some detail by Odling-Smee, Laland, and Feldman (Odling-Smee 1988; Laland et al. 1996, 1999, 2001). This initial work gave rise to several theoretical studies exploring different aspects of NC (reviewed in Odling-Smee et al. 2013), such as the effects of metapopulation structure on evolutionary dynamics of NC-related traits (Hui and Yue 2005; Borenstein et al. 2006; Silver and Di Paolo 2006; Han et al. 2006), inclusive fitness analyses of the evolutionary dynamics of NC-related traits (Lehmann 2008), the consequences of NC for kin selection and the evolution of co-operation (Lehmann 2007; Van Dyken and Wade 2012; Connelly et al. 2016), effects of niche construction on metapopulation dynamics (Hui et al. 2004; Han et al. 2009; Zhang F. et al. 2012), and multi-strain or multi-species population dynamics (Krakauer et al. 2009).

Before we proceed to our critique, let us briefly state our position on NC right at the outset. We do not doubt that NC, even in a meaningfully narrower sense than used by its proponents (e.g. as cogently delineated by Dawkins 2004; Brodie III 2005), is an important and reasonably common ecological phenomenon that can have interesting evolutionary consequences. We disagree, however, that the phenomenon has been ‘neglected’ in SET, and we will argue that, on the contrary, the phenomenon has been extensively incorporated in both ecological and evolutionary studies for at least over a century. We also disagree with the quasi-philosophical arguments of its proponents that the NC perspective “entails that niche construction be regarded as a fundamental evolutionary process in its own right” (Laland et al. 2016), and we demonstrate the fallacies and misconceptions that underlie this assertion. We trace some of these misconceptions to fundamental confusions among the proponents of NCT about what the conceptual core of SET actually is: they appear to conflate a narrow one-locus population genetic representation of the evolutionary process with evolutionary theory. In particular, the proponents of NCT seem to be completely unaware of, or at least not engaged with, quantitative genetic thinking in evolutionary theory. Finally, we show that much of the published literature on NCT is not only unnecessarily repetitive, and muddled biologically and philosophically, but is also historically inaccurate in trying to claim a degree of originality for NCT that it just does not have. Overall, this leads us to suspect that NCT is less a serious and consequential evolutionary theory and more an example of academic niche construction in a nascent post-truth scientific world.

### Niche construction has not been ‘neglected’ in SET

The proponents of NCT insist, despite criticism, on a very wide, all-encompassing definition of NC (Laland et al. 2005). Consider, for example, the following quotes from the NCT canon: “Organisms, through their metabolism, their activities, and their choices, define, partly create, and partly destroy their own niches. We refer to these phenomena as niche construction” (Odling-Smee et al. 1996) and “Niche construction is the process whereby organisms, through their metabolism, activities, and choices, modify their own and/or each others’ niches” (Odling-Smee et al. 2003). Given that everything an organism does, including living or dying, affects the environment, NC would appear to be a synonym of biology, in which case, we fail to see how it could be “neglected”. Nevertheless, the claim that NC has been “neglected” in SET is repeatedly made by the proponents of NCT, including in the eponymous sub-title of their book “*Niche construction: the neglected process in evolution*” (Odling-Smee et al. 2003). Even if we take a narrower definition of NC, as opposed to “niche changing”, as suggested by Dawkins (2004) and Brodie III (2005), it is hard to agree with the claim of “neglect”. Dawkins (2004) and Brodie III (2005), very correctly in our opinion, focus on the crucial issue of whether there are covariances between the variants of a ‘constructed’ aspect of the environment, variants of heritable organismal phenotypes, and the varying fitnesses of the latter. They argue that, if such covariances exist, then the phenomenon should be considered niche construction and, if they do not, it should be labelled niche changing, to emphasize that the environmental modification is simply a by-product, not the adaptive consequence of a variant phenotype’s activity (Dawkins 2004; Brodie III 2005). Even with this narrower definition of NC, the argument that SET has typically avoided incorporating a perspective wherein organisms can shape selection pressures, for themselves and <u>f</u>or other species, by altering the environment does not really stand in the face of the evidence, as we shall show below.

The core of the Darwinian view of evolutionary change is the notion that the ecological struggle for existence can result in evolutionary change because heredity (in the sense of parent-offspring similarity) mediates the greater representation of ecologically successful variants in subsequent generations (Dobzhansky 1937; Gayon 1998). The struggle for existence is itself a very Malthusian (Malthus 1798) metaphor, premised upon the fact that increased depletion of available resources by an increasing population eventually has a negative impact upon population growth rates and, consequently, selects for greater competitive ability. Thus, individuals alter their environment as a result of feeding and, consequently, affect the selection pressures faced by themselves: in other words, a classic case of what is now labelled NC, but lurking at the very heart of the foundations of SET. If NC is enshrined at the core of SET, it is a bit odd to find repeated claims in the NCT literature of its neglect by SET and its practitioners. Indeed, it is our submission that phenomena that are now sought to be highlighted under the label of NC have been extensively incorporated into explanations of various ecological and evolutionary processes for well over the last 100 years. In the fields of population and community ecology, for example, the incorporation of such NC phenomena began with models of density-dependent population regulation like the logistic (Quetelet 1835), alluded to above. Subsequently, an NC perspective continued to be incorporated into experimental and manipulative studies aimed at elucidating the biological mechanisms of density-dependent population growth regulation and community structuring via competition or predation (reviewed in Kingsland 1982).

Another striking example of an NC perspective at the very core of SET is to be found in Fisher’s (1918)conceptualization of the rest of the genome, including its allelic homologue, as constituting part of the environment of a focal allele at a given locus (Edwards 2014). Indeed, Fisher (1941) explicitly recognized that evolutionary change of allele frequency of the focal allele due to selection typically led to a change in the environment, including the ‘genomic environment’, in a manner that altered fitness of and, therefore, selection pressures on, the focal allele (discussed in Frank 1995). For example, consider the simple case of a one-locus, two-allele, viability selection model with over-dominance for fitness (i.e. *ω*_12_> *ω*_11_,*ω*_22_ where *ω*_11_, *ω*_12_ and *ω*_22_ are the fitnesses of the *A*_1_*A*_1_, *A*_1_*A*_2_ and*A*_2_*A*_2_ genotypes, respectively). Here, the steady state is a stable equilibrium allele frequency. Suppose the allele frequency of allele *A*_1_, say *p*
_1_, is less than the equilibrium value 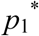
. In that case, in the next generation, *p*_1_ will increase. This increase automatically reduces the frequency of the allele *A*_2_ (*p*_2_ = 1- *p*_1_), which in Fisher’s (1918, Fisher’s 1941) view is an alteration of the environment, resulting in a reduction of the marginal allelic fitness of *A*_1_ (marginal allelic fitness of *A*_1_= *p*_1_*ω*_11_ + *p*_2_
*ω*_12_). Thus, the very increase of *p*_1_ as a result of selection (differential fitnesses of the three genotypes) in itself affects the genomic environment at that locus and results in a reduction of the rate of increase of *p*_1_. This is an excellent example of the kind of nuanced thinking about organism (in this case, allele) to environment feedbacks, resulting in an alteration of selection pressures, that NCT proponents claim is lacking in SET and, once again, it is found at the very heart of “gene-centric” SET.

Shifting our attention to theoretical studies of slightly more ecologically rich phenomena in evolution, like speciation, we again find a nuanced NC-like perspective even in studies firmly within the gene-centric SET framework, conducted by people who would have defined themselves as firmly within the SET tradition. Consider the class of population genetic models of sympatric speciation, which began with Maynard Smith’s (1966) eponymous paper, and are reviewed by Gavrilets (2006). In these models, the fitnesses of genotypes at a locus vary depending on which sub-habitat or niche is chosen by an organism. Such choice of habitat, incidentally, is considered to be NC by its proponents (Odling-Smee et al. 2003). In this set of models, choice of sub-habitat is arbitrarily fixed as a parameter and not modelled as being determined by genotypes at a locus, because the focus of the models is on the evolution of mating preferences. Similar models of host-choice evolution and sympatric speciation (e.g. Rausher 1984; Diehl and Bush 1989) have also been developed in which host choice is determined by genotypes at a preference locus, with the choice of host then determining fitnesses based on genotypes at a performance locus. Treating fitnesses on hosts 1 and 2 as conceptually the same as their being two separate traits (Falconer 1960), if there is antagonistic pleiotropy for fitness on alternative hosts, it drives epistasis for fitness between the preference and performance loci. If host preference in these models is specifically for oviposition, then the epistasis is trans-generational between maternal preference genotype and offspring performance genotype. All these models, thus, incorporate a very nuanced treatment of organisms affecting the selection pressures they (or their offspring) face through their choice of host or sub-habitat, something that the proponents of NCT maintain is generally missing in studies done within the SET framework.

If we switch our focus to empirical studies, we find that here, too, what is today called the NC perspective has actually been quite pervasive. In the relatively simplistic context of controlled-environment, single-species experimental evolution studies, experimenters impose selection pressures on laboratory populations and examine how populations evolve in response to them (Garland and Rose 2009). Even in such simple laboratory systems, nuanced perspectives, that today would be labelled NC, have been deployed to understand evolutionary phenomena, long before NC became a fashionable appellation (e.g. see Mueller et al. 2005 for a review). For example, in a study of the evolution of adaptations to larval crowding in *Drosophila melanogaster*, Borash et al. (1998) detailed how the reducing food and increasing nitrogenous waste levels in the deteriorating environment of a crowded culture vial resulted in temporally varying selection pressures within a generation. This pattern of changing selection pressures within a generation, in turn, mediated the evolution of a polymorphism, with early-developing larvae being faster feeders and late-developing larvae being more waste tolerant. These temporal changes in the environment, and the selection pressures they affected, were directly caused by the activities of the feeding and excreting larvae. Subsequently, similar studies have revealed that the evolutionary outcomes of the selection pressures resulting from the deterioration of the food environment in a crowded *Drosophila* culture further depend on aspects of the environment, such as the total amount of food available for waste to diffuse into (Nagarajan et al. 2016; Sarangi et al. 2016), and whether or not there are opportunities, resulting fortuitously from the husbandry techniques employed, for assortative mating with regard to development time (Archana 2010). The point we wish to stress is that the authors of all these studies have undertaken a very nuanced approach to understanding the selection pressures facing populations of *Drosophila* in cultures subjected to larval crowding in various ways. Their approach includes an explicit incorporation of the manner in which the activities of the larvae alter the environment and how that, in turn, modifies the selection pressures they face. These selection pressures then, in turn, interact with other specific aspects of the environment to result in varying evolutionary each others’ trajectories across studies. The results of many such studies linking demography, life-history, adaptations to crowding and population dynamics are reviewed by Mueller et al. (2005). We emphasize here that all these authors would place themselves squarely within the SET tradition and they have accomplished their analyses of a complex interplay between organism and environment in an evolutionary context without recourse to the specific conceptualizations or terminology of NCT. These examples, to our minds, are particularly striking because these are studies conducted within the laboratory selection paradigm, a scenario most likely to approximate the criticism of neglecting organism-to-environment-to-selection feedback that is routinely levelled against practitioners of SET by the proponents of NCT.

If one shifts focus from laboratory selection studies to studies of evolutionary phenomena in wild populations, the NC perspective is even more pervasive, although without the use of the NC label. As Thompson (1994) has detailed, a nuanced and detailed appreciation of how interacting organisms shape each other’s selection pressures and evolution is apparent even in the earliest late-nineteenth century studies of flower-pollinator coevolution and mimicry, some of the first attempts to understand diversity through the lens of natural selection. Indeed, the vast number of studies in the broad area of coevolution are suffused with the kind of appreciation of how subtle the reciprocal interactions between organisms and their biotic and abiotic environments are, and how they shape coevolutionary trajectories, that NCT proponents claim is generally missing in studies within the framework of SET. These studies encompass varied ecological and evolutionary nuances of species interactions ranging from competition to mutualism, including various aspects of predation, grazing and parasitism. This huge body of work is discussed in detail by Thompson (1994), 2005, 2013) and we will not dwell on it further.

Even in the realm of human cultural (or gene-culture) evolution, which is often highlighted as a clinching example of how a NC perspective yields insights beyond those available using SET (Laland et al. 2000, 2001, 2016; Odling-Smee et al. 2003; Laland and Sterelny 2006; O’Brien and Laland 2012), the claims of NCT proponents are somewhat overblown. The two examples most commonly cited in this context by NCT proponents are those of the evolution of the ability to use lactose as adults in human societies with a history of cattle husbandry and the high incidence of sickle cell anaemia in human populations with a high incidence of malaria due to the adoption of yam cultivation, respectively (Laland et al. 2000, Laland et al. 2001; Odling-Smee et al. 2003; Laland and Sterelny 2006; O'Brien and Laland 2012). However, our understanding of the causes underlying these evolutionary phenomena, and their rooting in human animal husbandry or farming activities, precedes the development of NCT by one or two decades (Wallach 2016). Indeed, to quote Wallach (2016): “These phenomena, as well as their causal relations to human activities, were not predicted or inferred from NCT, *nor was the formulation of NCT required for their explanation* (emphasis in the original)” and “In both cases, it is gene-culture co-evolution that does all the explanatory work, with NCT contributing nothing but an - arguably, apt - descriptive metaphor”.

Another claim by the proponents of NCT that is connected with the “neglect” argument is that Odling-Smee (1988) was the “first to introduce the concept of ‘ecological inheritance’” (Laland et al. 2016). This claim, too, is untenable, together with similar claims about how SET has neglected phenomena like phenotypic plasticity and NC (Laland et al. 2014c). Almost from its inception, quantitative genetics has been concerned with what are now called ecological inheritance and phenotypic plasticity (discussed in Prasad et al. 2015). Even basic textbooks of quantitative genetics (e.g. Falconer 1960) include extensive discussion of phenotypic similarity between parents and offspring due to shared environments, resulting in a positive parent-offspring covariance for environmental effects on phenotypic value. This is exactly the same concept that is now termed ecological inheritance in NCT. The notion in quantitative genetics of a covariance between genotypic value and environmental value of a trait clearly reflects a point of view that at least clearly acknowledges that genotype can affect the manner in which environment affects phenotype. This could, of course, result from certain genotypes responding differentially to certain environments but, equally, it is clear that genotype-environment covariance also encompasses all phenomena that are labelled as NC. Similarly, the very partitioning of phenotypic value into a genotypic and an environmental component in quantitative genetics exemplifies an appreciation of phenotypic plasticity as a concept and a phenomenon. Moreover, the quantitative genetic notion of genotype × environment interaction reflects an appreciation that there may be genetic variation for the degree and nature of phenotypic plasticity in a population. Indeed, the recent deployment of the concept of inclusive heritability (Danchin and Wagner 2010) in the context of an expanded and slightly modified quantitative genetics framework for analysing evolutionary change (Danchin et al. 2011), just serves to underscore the versatility and flexibility of the original, essentially phenotypic, formulation of evolutionary change under selection in classical quantitative genetics. While the proponents of NCT cite some of this work (e.g. in Laland et al. 2016), they seem to entirely miss the important corollary that, if all forms of non-genic inheritance fit comfortably within the basic framework of quantitative genetics, then clearly the classical formulations of SET are far more flexible and encompassing than the proponents of NCT give them credit for.

To sum up, we believe that the claim by proponents of NCT that NC and ecological inheritance have been neglected in evolutionary biology research carried out under the framework of SET is not tenable. On the contrary, the kind of nuanced appreciation of how organisms shape their abiotic and biotic environments and, thereby, the selection pressures they face, which the proponents of NCT claim to be largely missing from work done in the SET framework, is actually ubiquitous in such work. Such a nuanced appreciation of the interactive nature of ecological relationships is abundantly reflected in mainstream evolutionary biological work right from the time of Darwin and Haeckel (Gayon 1998; Richards 2008), through the decades of the crystallization of SET in the early twentieth century (Gayon 1998), and well into present times (Thompson 1994, Thompson 2005, Thompson 2013). Moreover, this nuanced appreciation of the richness of ecological relationships, and its deployment in evolutionary explanation, is seen in *a priori* ‘gene-centric’ theory, in theory slightly more responsive to ecological realities, whether in the laboratory or the wild, and in empirical studies on laboratory or wild populations and communities. Given all the examples quoted above, and there are countless more for each claim that we could not quote for want of space, our question to the proponents of NCT is: What *exactly* has been neglected by SET that NCT claims to incorporate?

### NCT models do not constitute an ‘extensive body of formal theory'

Here, we first briefly discuss three early and often cited models of niche construction that have been described as extending the understanding possible through SET and also as constituting an extensive body of formal theory (Laland et al. 2016) (these models are further discussed at length in Appendix 1). We then touch upon some other theoretical developments in NCT. Two of the three early models (Laland et al. 1996, 1999) are based on standard di-allelic two locus population genetic models with multiplicative fitnesses (general discussion in Hartl and Clark 1989), where one locus specifies a niche constructing phenotype which, in turn, affects fitness through genotypes at the second locus as a result of the specific environmental perturbations it causes. The third model (Laland et al. 2001) is a gene-culture coevolution model where the niche constructing trait is culturally inherited. Our main purpose in presenting the models in the appendix is to show the mathematically inclined readers how these models are simple extensions/variants of standard population genetics formulations. The mathematically disinterested reader can safely ignore Appendix 1 and continue reading about our major comments on these models.

These three early models consider a very specific form of niche construction, i.e., where the niche constructing activity of individuals, mediated by locus or cultural trait *E*, affects a common resource utilized by all members of the population. This resource affects selection at another locus *A* and the selection pressure, thus, depends on the frequency of allele/trait *E*. Consequently, the benefits or costs of niche construction are shared by all individuals irrespective of their own niche construction ability. Thus, these models cannot be used to analyse the effects or niche constructing phenotypes such as nest building, parental care, or habitat selection. Laland et al. (2005) have conceded this point, raised by Okasha (2005), and agree that a lack of spatial structuring in their models allows all individuals to benefit from the effects of niche construction, irrespective of whether or not they express the niche constructing phenotype.

The results from this set of models are what one would expect from a two-locus multiplicative fitness model with fitness at one locus being affected by the other in an epistatic and frequency-dependent manner, even if there were no niche construction involved (e.g. Bodmer and Felsenstein 1967; Karlin and Feldman 1970; Feldman et al. 1975; Karlin and Liberman 1979; Hastings 1981; Christiansen 1990). These results include changing the position of equilibria expected in a situation without frequency-dependent selection, appearance of new polymorphisms, disappearance of otherwise expected polymorphisms, fixation of traits which are otherwise deleterious, etc. (for details, see Appendix 1). The point at which niche construction differentiates these models from the ones without niche construction is the appearance of time lags between selection on the niche constructing locus and the response to changes in the allele/trait frequencies at the other locus. This can also lead to evolutionary momentum where selection at the second locus continues for some time after selection at the niche constructing locus has stopped or reversed. Similar effects have been seen in population genetic models of host specialization where there is a generational lag in the epistatic effects on fitness of genotypes at a maternal preference locus and an offspring performance locus (Rausher 1984; Diehl and Bush 1989), as well as in models of maternal inheritance (Kirkpatrick and Lande 1989) and early, pre-NCT, gene-culture coevolution (Feldman and Cavalli-Sforza 1976).

The main point we wish to make is that the results of these models are neither surprising nor unexpected in the context of SET. Two (or more) locus models with pleiotropic and epistatic effects on fitness, with or without time lags, are well known within gene-centric SET to yield outcomes and dynamics that can differ from simpler, similar models without epistasis (Hartl and Clark 1989). Indeed, gene-by-gene interactions, whether within- or between-loci, can yield seemingly unexpected results, even without time-lags. For example, under-dominance for fitness at one locus can, seemingly paradoxically, result in the fixation of a sub-optimal phenotype due to initial conditions fortuitously being what they were, unlike what is possible in a one-locus model of viability selection without under-dominance. We do not believe that it therefore follows that under-dominance is a major evolutionary process in its own right, or that a few models of over-dominance and under-dominance need to be elevated to the status of a major evolutionary theory to be set up in competition to SET. Essentially, the ‘formal theory’ pertaining to models of the evolutionary consequences of niche construction rests upon the demonstration that, because niche construction can induce time-lagged epistasis for fitness, it can result in evolutionary outcomes that are unexpected in the context of simpler population genetic models lacking such time-lagged epistatic effects. This does not in any way supersede SET, unless one implicitly defines SET as taking no cognizance of gene-by-gene interactions.

In addition to the three models we discuss in the appendix (Laland et al. 1996, 1999, 2001), the “extensive body of formal theory (that) explores the evolutionary consequences of niche construction and its ramifications for evolutionary biology and ecology” (Laland et al. 2016) consists of a handful of models, some of which extend the logic of Laland et al. (2001) by showing how incorporating NC into simpler models can result in novel outcomes, including the evolution of niche construction (Silver and Di Paolo 2006; Creanza and Feldman 2014). Some other models explore the role of NC in facilitating altruism (Lehmann 2008; Van Dyken and Wade 2012), whereas others suggest that NC can facilitate range expansion and coevolution (Kylafis and Loreau 2008; Krakauer et al. 2009), neither of which are a great surprise to practitioners of SET. We frankly fail to understand how or why we are expected to acknowledge this handful of models as an extensive body of formal theory that somehow significantly supersedes or, at least significantly extends, SET. These papers taken together would not even compare favourably with one SET paper, that of Fisher (1918), in terms of the novel evolutionary insights they provide.

In light of the arguments above, and the detailed discussion in Appendix 1, we pose two questions here. First, what are the theoretical papers on NCT (barring those that we cite ourselves) that, taken together, could justify the label “extensive body of formal theory”? Second, what are the major theoretical insights emanating from this “extensive body of formal theory” that are not intuitively obvious from analogous SET formulations?

### NC is not an evolutionary process at par with natural selection

The proponents of NCT also make some quasi-philosophical claims that (a) NC and developmental bias are essentially two sides of the same coin, one internal and one external to the organism (Laland et al. 2008, 2014b); (b) NC is a fundamental evolutionary process in its own right (Odling-Smee et al. 2013); and (c) NC as an ‘evolutionary process’ is somehow logically at par with natural selection in terms of its importance to ‘causal’ evolutionary explanations of adaptation (Laland 2015). One somewhat unique characteristic of the literature on NC is that the same few conceptual arguments are repeatedly made, in very similar words, in multiple publications. Consequently, to avoid having to keep referencing multiple papers that say almost the same things, we focus our critique of these quasi-philosophical claims on their most recent detailed exposition (Laland 2015).

Essentially, these claims are premised upon the argument that the process of adaptive evolution has two major steps: the generation of variation and the sorting of this variation such that the frequency of better adapted variants increases. This is an old and venerable view, going back at least to Bateson (1894) and De Vries (1909), and also articulated in some detail in recent times by Endler (1986). We agree with this depiction of the adaptive evolutionary process. Laland (2015) argues that phenomena acting at the first of these two steps have not typically been regarded as ‘evolutionary processes’, whereas phenomena acting at the second step, such as natural selection, have. Laland (2015) further traces the roots of this distinction between evolutionary processes like selection on the one hand, and background conditions that alter the form of selection, or the extent of variation available to selection, on the other, to Mayr’s (1961) distinction between proximate and ultimate causes in biological explanation. Specifically, Laland (2015) argues that background conditions, such as niche construction or developmental bias, that shape the specific instantiation of a (proximal) causal process like selection should not be neglected as causal processes. We believe that Laland (2015) misses the point that natural selection is an ultimate cause when thinking of a phenotype, but a proximate cause when thinking of change in the composition of a population with regard to the variants it contains. For example, if one is interested in the differences in beak shape among the various species of Darwin’s finches on the Galapagos islands, natural selection is invoked as an ultimate cause, whereas the proximate cause(s) are to be sought in the ontogeny of beak development in the different species. However, if one is trying to understand why average beak length in a given species of Darwin’s finch changes due to, say, a drought, then a specific instance of natural selection constitutes a proximate cause. Thus, the proximate-ultimate distinction in this context is really a red herring, because whether natural selection is a proximate or an ultimate cause depends upon what exactly is sought to be explained. Essentially, natural selection is as much a proximate cause of evolutionary change as developmental-physiological mechanisms are proximate causes of a phenotype. It is not clear to us why the proponents of NCT consider proximate causes to be very important when thinking of phenotypes, but not when thinking of evolutionary change as the phenomenon whose various causes are to be delineated. Indeed, the relevant literature by NCT proponents gets fairly muddled on this point. The real issue is actually the contrast between a phenomenon acting at the ‘sorting of variation’ stage and one acting at the ‘generation of variation’ stage.

As stated above, Laland (2015) argues that background conditions that shape the specific instantiation of a (proximal) causal process like selection should not themselves be neglected as causal processes. In a very broad sense, we do not disagree that background conditions can affect the nature of the outcome of a proximate causal process and, in that sense, considering background conditions can add to the richness of an explanation. However, this added richness is often of a specific and narrow kind. We will elaborate upon this point using Laland’s (2015) analogy of a murder trial in which he writes, “we would not be optimistic about the chances of the defendant in the dock receiving a ‘not guilty’ verdict if their defence was based on the argument that they did not cause the death of the victim that they shot – that was the bullet – they only pulled the trigger”. In this analogy, the evolutionary change is the death of the victim, the bullet is natural selection, and the defendant deciding to pull the trigger is the background condition shaping the specific form that selection took in this particular instantiation. The argument deployed by Laland (2015) is that the judge, presumably representing an enlightened NC theorist, will give more, or equal, importance to the background condition as compared to selection. However, the analogy is actually flawed because the judge in this example is specifically tasked with ascertaining human agency – his or her brief is to ascertain whether the defendant is guilty of the murder or not. In the mapping of this analogy on to the evolutionary process, there is no equivalent of the need to establish agency. Evolutionary theory attempts an explanation of the process of adaptive evolution, and steers clear of any requirement of establishing agency for one or another process. Once the consideration of establishing agency is removed, and the focus is on the facts of the case, as would be the situation for evolutionary theory, the situation becomes quite different from that presented by Laland (2015): an autopsy report, which is not concerned with agency and is therefore the appropriate analogue of evolutionary theory in this example, will simply record the cause of death as a bullet wound suffered by the victim!

The error implicit in conflating the logical causal status of proximate causal processes and background conditions can be seen clearly using a different analogy, also used to great effect by Darwin (quoted at length in Gayon 1998, pgs. 52-53). The designing and construction of a building is constrained first by the laws of physics and, secondarily, by the availability of materials due to accessibility or finances or both. But within these background constraints, architects and their style of architecture exert great influence on the final form that the building takes. Laland’s (2015) argument is essentially a plea for giving equal importance to the laws of physics and material availability on the one hand, and the design of the architect on the other, as an explanation of the final form of a building. While not denying the role of the background conditions, we would suggest that the proximate cause of the architect’s design has far greater explanatory power in explaining the final form of the building, as compared to the background conditions. That is why, despite the laws of physics, the use of white marble as the primary construction material, and even the raison d’être, being the same in both cases, the Lincoln Memorial in Washington DC and the Taj Mahal in Agra do not look particularly similar: the differences arise from one being designed in the Doric style and the other in the Indo-Persian style.

The second aspect of Laland’s (2015) conflation of NC and selection as causal evolutionary processes that we disagree with stems from our distinction between a phenomenon *per se* and the conditions that cause the phenomenon to occur in a particular manner in a specific instantiation of that phenomenon. Natural selection is a phenomenon that results from the sorting of variants such that the frequency of better adapted variants increases. NC is a phenomenon that can shape the manner in which selection acts in any particular case and, thereby, affect precisely which variants end up increasing or decreasing in frequency as a result of selection. These two phenomena are clearly not of the same type with regard to their roles in the adaptive evolutionary process. NC affects the way in which selection acts. Its role is thus of a modifier which affects how a certain category of evolutionary process acts in a given instantiation, whereas selection has a very different logical or epistemic status as a specific category of process. We see the attempts to join NC with developmental bias (Laland et al. 2008, 2014b; Laland 2015, 16) as an essentially flawed attempt to make this joint phenomenon appear more important than NC alone, because conflating the two also gives the joint phenomenon a double role in affecting the ‘generation of variation’ step as well as mediating the specific instantiation of selection in the ‘sorting of variation’ step. Nevertheless, it does not alter the fact that it both cases, it is a specific instantiation of selection that is being altered by NC or development bias.

To sum up, although we agree that both the generation and sorting of variants are important parts of the overall process of adaptive evolution, we do not agree that NC/developmental bias have a logical or epistemic status in explanations of adaptive evolution equivalent to that of natural selection. In the context of adaptive evolutionary change, natural selection is the proximate cause of changes in population composition and is, therefore, a general principle rather than a phenomenon that exists as a specific instantiation of a general principle. Consequently, the position or status of natural selection in explanations of adaptive evolution is epistemically distinct from that of phenomena that either constrain the range of variation that selection sorts, or that modulate which variants selection happens to sort for in a given scenario.

## Concluding remarks

Over the past decade or so, there has been an increasing realization that there are serious problems with the way science is being done, published and evaluated (Lawrence 2003; Ioannidis 2005; Song et al. 2010; American Society for Cell Biology 2012; Balaram 2013; Head et al. 2015; Horton 2015; Smaldino and McElreath 2016). Of particular concern (possibly even replacing plagiarism concerns of a few years prior), are an increasing tendency to not reference prior work properly, in order to be able to bolster exaggerated claims of novelty (Robinson and Goodman 2011; Teixeira et al. 2013; Maes 2015), as well as an increasing trend of setting up one’s work as a competing counterpoint to some dominant idea in the field, even if the work is actually complementary to that dominant idea. The published work on NCT, especially after Odling-Smee et al. (2003), exemplifies this state of affairs.

Proponents of NCT make a few claims repeatedly: (i) NC and ecological inheritance have been neglected; (ii) there is a vast body of formal theory on NC and its ecological and evolutionary consequences that is a significant addition to SET; and (iii) NC and, more recently, NC/developmental bias are important evolutionary processes at par with natural selection in the context of explaining adaptive evolution. These claims are repeated, often in similar language and with the same examples and analogies, in paper after paper. Indeed, examining the fairly voluminous literature on NCT after the book by Odling-Smee et al. (2003), we find that hardly anything new has been said, with the exception of the more recent claims that developmental bias and niche construction are conceptually two sides of the same coin (Laland et al. 2008; Laland et al. 2014b; Laland 2015, 2016). Characteristically, even this claim has been repeated multiple times over the past several years, without any new arguments or facts being deployed to bolster it.

As we have argued here, we believe the facts clearly suggest that the first two claims are just plain wrong. We wonder whether that is why they need so much repetition. The third claim, we have argued here, is based on philosophically muddled thinking and inappropriate analogies. Thus, we believe that the scientific case being made by the proponents of NCT is weak, and their work and perspective is being sought to be made to appear far more novel, revolutionary and consequential than it really is. It is this latter point that we find disturbing and, more importantly, detrimental to the way in which science is done. Incessant repetition of claims that do not stand up to critical scrutiny, an avoidance of specifically responding to particular criticisms in favour of diffuse and generalized responses, and the deployment of dubious analogies are all aspects of rhetoric that are unfortunately becoming familiar worldwide in what is often being described as a post-truth world. We are dismayed that they have also made an entry into the scientific discourse and that is why we wonder whether this constant pushing of untenable claims regarding NCT is actually an instantiation of academic niche construction.

## Acknowledgments

This is contribution no. 2 from the Foundations of Genetics and Evolution Group (FOGEG). FOGEG is an informal association of SD, AJ, NGP and TNCV (and, sometimes, some of their students) getting together periodically to work on conceptual issues at the foundations of genetics and evolutionary biology. All four FOGEG authors contribute equally to the manuscripts and the sequence of these four authors is chosen at random for each submission. The last author acts as corresponding author for that submission. AJ thanks the Department of Science and Technology, Government of India, for support via a J. C. Bose Fellowship. SD, NGP and TNCV thank IISER Pune, IISER Mohali, and JNCASR, respectively, for in-house funding. MG is supported by a scholarship from JNCASR. We also thank the anonymous reviewers for helpful comments on the manuscript.

## Appendix 1: Models of Niche Construction

Here we discuss, in some detail, the three early and often cited models of niche construction that have been described as extending the understanding possible through SET, and as constituting an extensive body of formal theory (Laland et al. 2016). Two of these models (Laland et al. 1996, 1999) are based on standard di-allelic two locus population genetic models with multiplicative fitnesses (general discussion in Hartl and Clark 1989), where one locus specifies a niche constructing phenotype which, in turn, affects fitness through genotypes at the second locus as a result of the specific environmental perturbations it causes. The third model (Laland et al. 2001) is a gene-culture coevolution model where the niche constructing trait is culturally inherited. We start by describing the di-allelic two locus genetic models (Laland et al. 1996, 1999). This appendix is aimed at readers familiar with population genetics models in order to show clearly how these foundational NCT models do not add anything substantial to what is already known from standard two-locus viability selection models incorporating frequency-dependent and epistatic effects on fitness.

**Genetic models** In these models, the niche constructing activity is controlled by a locus labelled *E*. At this locus, there are two alleles, *E* and *e*, and the niche constructing ability of the population, reflected entirely by resource levels, is directly proportional to the frequency of the allele *E*, given by *p*. A resource, *R*, is defined such that its amount is directly proportional to the niche constructing activities of the present and past generations. In the first model (Laland et al. 1996), *R* depends on *n* previous generations of niche construction and the *n* generations can either weigh in equally, or there can be recency or primacy effects. A recency effect entails generations closer to the present generation having a larger effect on *R* than the ones further in the past. A primacy effect entails generations further into the past having a larger effect on *R* than the ones which are more recent. In the second model (Laland et al. 1999), *R* is also affected by autonomous ecological mechanisms of resource depletion or recovery, besides the niche constructing activity of the population. In this case, *R* is given by the following recursion equation:

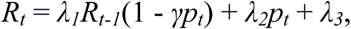

where *R_t_* is the amount of resource in the present generation, *R_
t-1_* is the amount of resource in the previous generation, *λ_1_* determines independent depletion, *λ_2_* determines effect of positive niche construction, *λ_3_* determines independent renewal, and *γ* determines effect of negative niche construction. In both models, *R* is constrained to be between 0 and 1 (0<*R*<1) and the value of *R* determines fitness through genotypes at a second locus, *A*, with alleles, *A* and *a*. *A* is favoured when *R* is high (above 0.5), whereas *a* is favoured when *R* is low (below 0.5). The two-locus genotypic fitnesses are given in Table 1 in terms of *R* and marginal one locus genotypic fitnesses.

**Table 1.** Matrix of two-locus genotypic fitnesses for the first two models (Laland et al. 1996, Laland et al. 1999). The symbols in brackets alongside the one locus genotype symbols are one locus marginal genotypic fitnesses. *f* is either 0.5, 1, or 2 in the first model, whereas it is 1 in the second model.

| Locus | $EE (\alpha_1)$ | $Ee (1)$ | $Ee (\beta_1)$ |
| --- | --- | --- | --- |
| $AA (\alpha_2)$ | $w_{11} = \alpha_1 \alpha_2 + \varepsilon R^f$ | $w_{12} = \alpha_2 + \varepsilon R^f$ | $w_{13} = \beta_1 \alpha_2 + \varepsilon R^f$ |
| $Aa (1)$ | $w_{21} = \alpha_1 + \varepsilon \sqrt{(R(1-R))^f}$ | $w_{22} = 1 + \varepsilon \sqrt{(R(1-R))^f}$ | $w_{23} = \beta_1 + \varepsilon \sqrt{(R(1-R))^f}$ |
| $aa (\beta_2)$ | $w_{31} = \alpha_1\beta_2 + \varepsilon(1-R)^f$ | $W_{32} = \beta_2 + \varepsilon(1-R)^f$ | $w_{33} = \beta_1\beta_2 + \varepsilon(1-R)^f$ |

In Table 1, *ε* gives the strength and direction of niche construction (-1<*ε*<1). The two locus gametic frequencies are given by *x_1_*, *x_2_*, *x_3_*, *x_3_*, and *x_4_* for *EA*, *Ea*, *eA*, and *ea*, respectively. The recombination rate is given by *r*. Then, the gametic recursions are given by,

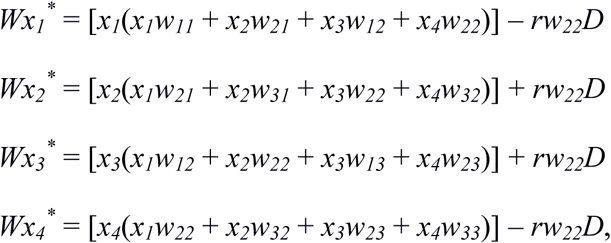

where 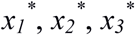
, and 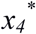
 are two locus gametic frequencies in the next generation, and *D* is the linkage disequilibrium given by,

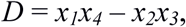

and *W*, the mean fitness of the population, is the sum of all the right-hand sides in the gametic recursions. The dynamics of this model were studied under four conditions, namely, no external selection at *E* or *A* loci, external selection at the *A* locus, external selection at the *E* locus, and overdominance at both loci.

*<u>1. No external selection</u>* No external selection means that frequency of the *E* allele remains constant (*α_1_* = *β_1_* = *α_2_* = *β_2_* = 1). Thus, the amount of resource remains constant in the first model and attains an equilibrium value in the second model. If these values are above 0.5, then allele *A* gets fixed and otherwise allele *a* gets fixed due to the selection generated by niche construction. The line, *R* = 0.5, defines a neutrally stable equilibrium in both models. Fixation of allele *A* is unstable below *R* = 0.5 and fixation of allele *a* is unstable above *R* = 0.5.

*<u>2. External selection at the E locus</u>* If selection favours allele *E* (*α_1_* > 1 > *β_1_*, *α_2_* = *β_2_* = 1), alleles *E* and *A* get fixed (*x_1_* = 1) if *ε* is positive and alleles *E* and *a* get fixed (*x_2_* = 1) if *ε* is negative. If selection favours allele *e* (*α_1_* < 1 < *β_1_*, *α_2_* = *β_2_* = 1), alleles *e* and *a* get fixed (*x_4_* = 1) if *ε* is positive and alleles *e* and *A* get fixed (*x_3_* = 1) if *ε* is negative. In the model with independent renewal and depletion of *R*, an additional caveat on the result is whether *R* is less than one half or more at the fixation value of *E* or *e*, respectively. Allele *A* gets fixed if *R* > 1/2 at fixation of either allele at the *E* locus, whereas allele *a* gets fixed if *R* < 1/2. A range of polymorphic equilibria are obtained if *R* = 1/2.

In the first model, if more than one previous generation of niche construction affects *R* (*n* > 1), then time lags between the start of selection at locus *E* and response at locus *A* can occur as *R* builds up slower than the rate of fixation at locus *E*. This evolutionary inertia is largest when there is a primacy effect and smallest when there is a recency effect. A similar process can lead to an evolutionary momentum type of effect as well when the selection at locus *E* stops or reverses because there is a lag between the frequency of allele *E* and the amount of resource accumulated. Such effects are not seen in the second model as there are no primacy effects in it. But, they can be obtained by making *λ_2_* = 1/*n*, i.e., a primacy effect.

In the second model, the rate of fixation of allele *A* when allele *E* is being favoured by external selection is dependent on magnitude of the impact of niche construction on the resource (*λ_2_*). Increasing the value of *λ_2_* increases the value of *R*, and, if *R* is near 1, this reduces the difference in fitness between the genotypes *AaEE* and *aaEE*, thus, remarkably reducing the rate of fixation of allele *A*.

*<u>3. External selection at the A locus</u>* In the first model, if external selection favours allele *A* (*α_1_* = *β_1_* = 1, *α*_*2*_ > 1 > *β_2_*) and there is no niche construction (*ε* = 0), or selection due to it is very weak (1- *α_2_ < ε* < 1- *β_2_*), allele *A* always gets fixed if it is present. Near *x_4_* = 1, if *ε* > 1- *β_2_*, fixation of allele *a* is neutrally stable. A set of polymorphic equilibria are possible near *x_1_* = 1, with alleles *e* and *a* increasing, if 1- *α_2_* > *ε*.

In the second model, if external selection favours allele *A* (*α_1_* = *β_1_* = 1, *α
_2_* > 1 > *β_2_*) and is weak, niche construction is positive (*λ*_*2*_ > 0, *γ* = 0), and *ε* is greater than zero, then for small values of *R* selection due to niche construction can take allele *a* to fixation. Also, there are a range of values of *R*, for which stable equilibria for fixation of alleles *A* and *a* overlap. If external selection is strong then fixation of allele *a* becomes less probable. If niche construction is negative (*λ_2_* = 0, *γ* >0 fixation of allele *a* still happens at low values of *R*, but, now those correspond to higher values of *p* instead of lower in the case of positive niche construction. For negative values of *ε* and positive niche construction, fixation of both *a* and *A* alleles becomes unstable and a set of stable polymorphisms are possible. If niche construction becomes negative, stable polymorphisms are possible near *x_3_* = 1 and allele *A* gets fixed for rest of the parameter space.

*<u>4. Overdominance at both loci</u>* Di-allelic two locus viability models can have a maximum of four gamete fixation states, four allelic fixation states, and seven interior fixation states (Karlin 1975). The results from overdominance (*α_1_*, *α_2_*, *β_3_*, *β_4_* < 1) are too complicated and varied to go into detail here, but generally, the effect of niche construction is to move the interior polymorphic equilibria and the edge equilibria (when they exist) towards higher values of *q*, when *R* is more than one half and towards lower values of *q* when *R* is less than one half. The magnitude of shift depends on the how far frequency of allele *E* is from one half. For high values of *ε* the edge equilibria can even merge with the respective gamete fixation states. For tightly linked loci (small *r*), niche construction can either increase or decrease linkage disequilibrium at genetic equilibrium. Equilibrium frequencies of allele *E* greater than one half (*p* > 1/2) result in increase in equilibrium frequencies of gametes *AE* and *Ae* and equilibrium frequencies of allele *E* less than one half (*p* > 1/2) result in decrease in equilibrium frequencies of gametes *aE* and *ae*. In the second model, these effects of niche construction persist, for some sets of parameter values, even when there is external renewal or depletion of the resource.

It is important to note that these two models have different meanings of positive and negative niche construction (Laland et al. 2005). For the first model, positive niche construction (*ε* > 0) means that increase in *R* increases the fitness of allele *A*. For the second model, positive niche construction implies that *λ_2_* > 0, *γ* = 0; negative niche construction implies that *λ_2_* = 0, *γ* > 0, meaning that increase in frequency of allele *E* (*p*) results in an increase in *R*, even though the sign of *ε* still mediates the effect of *R* on selection at the *A* locus.

***Cultural model*** We turn now to the third model in which the niche constructing trait is culturally inherited. A niche constructing trait *E* with variants *E* and *e* is postulated as a culturally inherited trait. A resource *R* depends on either *n* previous generations of niche construction, i.e., the frequency of trait variant *E*(*x*) (Model 1), or on niche construction and independent renewal or depletion following the same equation for *R* as in the second model (Model 2). A genetic locus *A* is postulated with alleles *A* and *a*, and its fitness is affected by amount of resource present with allele *A* being favoured when *R* > 1/2 and allele *a* being favoured when *R* < 1/2. The six pheno-genotypes, *AAE*, *AAe*, *AaE*, *Aae*, *aaE* and *aae*, have frequencies *z_
1_*-*z_6_* and their fitnesses are given in Table 2. Rules for vertical cultural transmission are given in Table 3.

**Table 2.** Matrix of pheno-genotypic fitnesses in terms of marginal trait/genotypic fitnesses given in brackets.

| | $E (\alpha_1)$ | $e (\alpha_2)$ |
| --- | --- | --- |
| $AA (\eta_1)$ | $w_{11} = \alpha_1 \eta_1 + \varepsilon R$ | $w_{12} = \alpha_2 \eta_1 + \varepsilon R$ |
| $Aa (1)$ | $w_{21} = \alpha_1 + \varepsilon \sqrt{R(1-R)}$ | $w_{22} = \alpha_2 + \varepsilon \sqrt{R(1-R)}$ |
| $aa (\eta_2)$ | $w_{31} = \alpha_1 \eta_2 + \varepsilon (1-R)$ | $w_{32} = \alpha_2 \eta_2 + \varepsilon (1-R)$ |

**Table 3.** Probabilities of offspring having trait *E* or *e* for each combination of parental mating.

|  | Offspring |  |
| --- | --- | --- |
| Matings | $E$ | $e$ |
| $E \times E$ | $b_3$ | $1 - b_3$ |
| $E \times e$ | $b_2$ | $1 - b_2$ |
| $e \times E$ | $b_1$ | $1 - b_1$ |
| $e \times e$ | $b_0$ | $1 - b_0$ |

Three specific cultural transmission scenarios were analysed: unbiased transmission (*b_3_* = 1, *b_2_* = *b_1_* = 0.5, *b
_0_* = 1), biased transmission (*b_3_* = 1, *b_2_* = *b_1_* = *b*, *b_0_* = 1, *b* ≠ 0.5), and incomplete transmission (*b_3_* = 1 - *δ*, *b_2_* = *b_1_* = *b*, *b_0_* = *δ*, *δ* > 0). For ease of analysis, the recursions were written in terms of allelo-phenotypic frequencies, namely, *AE*, *aE*, *Ae*, and *ae* (for the equations see Laland et al. 2001). Similar results were obtained for both model 1 and model 2 unless otherwise stated.

*<u>1. No external selection</u>* For unbiased transmission, the results for model 1 are analogous to Laland et al. (1996; see above) and the results for model 2 are analogous to Laland et al. (1999; see above).

For biased transmission, frequency of trait *E* increases if *b* > 0.5 and that of trait *e* increases if *b* < 0.5. For positive values of *ε* and *b* < 0.5, *ae* is fixed at equilibrium values of *R* < 0.5 and *Ae* is fixed at equilibrium values of *R* > 0.5. If *b* > 0.5, trait *E* gets fixed instead of trait *e*. Symmetric results are obtained when ε is negative.

For incomplete transmission, when *δ* > 0 and *b* = 0.5 the cultural trait remains polymorphic and a line of neutrally stable equilibria is obtained for locus *A*. If *b* ≠ 0 and *ε* is positive allele *A* gets fixed for equilibrium values of *R* > 0.5 and allele *a* get fixed for equilibrium values of *R* < 0.5. Symmetric results are obtained when *ε* is negative.

*<u>2. External selection at the A locus</u>* Again, for unbiased transmission, the results for model 1 are analogous to Laland et al. (1996; see above) and the results for model 2 are analogous to Laland et al. (1999; see above).

For biased transmission, when cultural transmission favours trait *E* (*b* > 0.5) and *ε* is positive, whether external selection at the *A* locus is opposed or not depends on the value of *R* at fixation of trait *E*. The positive *ε* and increasing frequency of trait *E* make it improbable that *R* will be lower than 0.5 at equilibrium. For cultural transmission favouring trait *e* (*b* < 0.5), *R* ends up being low enough for fixation of allele *a* instead of *A* more often. When *ε* is negative, three polymorphic equilibria are possible depending on value of *R*, namely, fixation of *AE*, fixation of *aE*, or an equilibrium polymorphic for alleles *A* and *a*. Symmetrically opposite results are obtained when niche construction is negative, i.e., trait *E* is responsible for depletion of the resource.

For incomplete transmission, a polymorphism for the cultural trait is obtained, and if *ε* is positive, either of the alleles *A* or *a* get fixed, depending on value of *R* at the equilibrium frequency of the cultural trait. If *ε* is negative, a fully polymorphic equilibrium is possible for very high values of *R* at equilibrium.

*<u>3. Selection at the cultural trait</u>* Again, for unbiased transmission, the results for model 1 are analogous to Laland et al. (1996; see above) and the results for model 2 are analogous to Laland et al. (1999; see above).

For biased transmission, when natural selection and transmission bias reinforce each other by either favouring *E* (*α_1_* > 1 > *α
_2_*; *b* > 0.5) or *e* (*α_1_* < 1 < *α_2_*; *b* < 0.5), *AE* or *ae* get fixed for positive values of *ε*. When these processes work against each other than their relative strength determines the final equilibrium. In such a scenario, cultural transmission can fix the trait which is not favoured by selection, if transmission bias is strong enough.

For incomplete transmission, the frequency of the cultural trait is given by a cubic equation (see Laland et al. 2001). For model 1, if *n* > 1 then time lags are obtained as in the analogous genetic model (Laland et al. 1996). The length of the time lag depends on both the selection coefficients and transmission bias with cultural transmission usually shortening the lags as compared to completely genetic models.

*<u>4. Overdominance at the A locus</u>* For unbiased transmission, polymorphisms at the locus *A* are possible if the selection due to niche construction does not completely overcome the external selection at the *A* locus, i.e., *R* is either too large or too small.

For biased transmission, polymorphisms at the *E* trait no longer exist and either trait *E* or e gets fixed. Frequency of alleles at the *A* locus depends on the interplay of external selection and selection due to niche construction.

For incomplete transmission, if there is no statistical association between the cultural trait and the genetic locus then a single polymorphic equilibrium is obtained. Selection due to niche construction shifts this equilibrium from the point where it would have been had niche construction not been acting.

